# Can genetic polymorphisms predict response variability to anodal transcranial direct current stimulation of the primary motor cortex?

**DOI:** 10.1101/2020.03.31.017798

**Authors:** Michael Pellegrini, Maryam Zoghi, Shapour Jaberzadeh

## Abstract

Genetic mediation of cortical plasticity and the role genetic variants play in previously observed response variability to transcranial direct current stimulation (tDCS) have become important issues in the tDCS literature in recent years. This study investigated whether inter-individual variability to tDCS was in-part genetically mediated. In sixty-one healthy males, anodal-tDCS (a-tDCS) and sham-tDCS were administered to the primary motor cortex at 1mA for 10-minutes via 6×4cm active and 7×5cm return electrodes. Twenty-five single-pulse transcranial magnetic stimulation (TMS) motor evoked potentials (MEP) were recorded to represent corticospinal excitability (CSE).

Twenty-five paired-pulse MEPs were recorded with 3ms inter-stimulus interval (ISI) to assess intracortical inhibition (ICI) via short-interval intracranial inhibition (SICI) and 10ms ISI for intracortical facilitation (ICF). Saliva samples tested for specific genetic polymorphisms in genes encoding for excitatory and inhibitory neuroreceptors. Individuals were sub-grouped based on a pre-determined threshold and via statistical cluster analysis. Two distinct subgroups were identified, increases in CSE following a-tDCS (i.e. Responders) and no increase or even reductions in CSE (i.e. Non-responders). No changes in ICI or ICF were reported. No relationships were reported between genetic polymorphisms in excitatory receptor genes and a-tDCS responders. An association was reported between a-tDCS responders and GABRA3 gene polymorphisms encoding for GABA-A receptors suggesting potential relationships between GABA-A receptor variations and capacity to undergo tDCS-induced cortical plasticity. In the largest tDCS study of its kind, this study presents an important step forward in determining the contribution genetic factors play in previously observed inter-individual variability to tDCS.

## Introduction

Transcranial direct current stimulation (tDCS), a popular non-invasive brain stimulation (NIBS) technique, alters the excitability of cortical regions by modulating the resting membrane potential of neurones underlying the electrodes. This causes changes in cortico-cortical excitability and corticospinal excitability (CSE) similar to the N-Methyl-D-aspartic acid receptor (NMDAR)-mediated mechanisms of long-term potentiation (LTP) and long-term depression (LTD) (M. A. Nitsche et al., 2003, 2007a). These changes are polarity dependent, anodal-tDCS (a-tDCS) depolarising the cortical area of interest, increasing CSE, and cathodal-tDCS (c-tDCS) hyperpolarises the cortical area of interest, reducing CSE (M. A. Nitsche & Paulus, 2000). This predictability has led to recent investigation into tDCS application in pathological population including Parkinson’s disease (Broeder et al., 2019; Bueno et al., 2019), epilepsy (Lin et al., 2018; Tecchio et al., 2018) stroke rehabilitation (Biou et al., 2019; Solomons & Shanmugasundaram, 2019), cerebral palsy (Inguaggiato et al., 2019), chronic pain (Callai et al., 2019; Jafarzadeh et al., 2019), depression (Aparicio et al., 2019) and cognitive impairment (Emonson et al., 2019).

Recent studies have challenged this predictability with systematic adjustments to initial standard tDCS protocols (M. A. Nitsche & Paulus, 2000) reporting findings opposing the historical predictability of tDCS. Increasing the stimulation duration of a-tDCS from the common 13-minutes to 26-minutes significantly reduced CSE, as measured by transcranial magnetic stimulation (TMS) evoked motor evoked potential (MEP) (Monte-Silva et al., 2013). Additionally, for c-tDCS, a reversal of the expected response was reported with significant increases in CSE, via TMS-evoked MEPs, when stimulus intensity of 20-minutes of c-tDCS increased from the common 1mA to 2mA (Batsikadze et al., 2013). These studies offer insight into divergences from traditionally expected responses to tDCS when adjusting stimulus parameters.

A growing number of published large sample-size studies have challenged the predictability of tDCS by reporting distinct subgroups of individuals either responding as historically expected to tDCS or not (Ammann et al., 2017; Chew et al., 2015; Labruna et al., 2016; López-Alonso et al., 2014, 2015; Puri et al., 2015, 2016; Strube et al., 2015, 2016; Tremblay et al., 2016; Wiethoff et al., 2014). Via a number of techniques, comprehensively reviewed elsewhere (Pellegrini et al., 2018b), participants have been sub-grouped into those who respond as historically expected to a-tDCS with increases in CSE (i.e. ‘responders’) and those who do not, responding with either no increase, or even reductions in CSE (i.e. ‘non-responders’). This dichotomy, termed inter-individual variability, extends to other NIBS protocols including Theta Burst Stimulation (TBS) (Goldsworthy et al., 2016; Hamada et al., 2013; Hinder et al., 2014; López-Alonso et al., 2014, 2015; Nettekoven et al., 2015; Puri et al., 2016; Vallence et al., 2015) and Paired Associative Stimulation (Labruna et al., 2016; López-Alonso et al., 2014, 2015; F. Müller-Dahlhaus et al., 2015; J. F. M. Müller-Dahlhaus et al., 2008; Strube et al., 2015). When keeping stimulus parameters consistent with standard protocol of 1mA current intensity and 10-13 minutes stimulus duration (M. A. Nitsche & Paulus, 2000), responder rates vary from 20-60% (Ammann et al., 2017; Chew et al., 2015; López-Alonso et al., 2014; Puri et al., 2016; Strube et al., 2015, 2016; Tremblay et al., 2016).

Several studies have recognised not all participating individuals are homogenous and intrinsic differences between individuals may be a source of inter-individual variability. Summarised elsewhere (Li et al., 2015; Pellegrini et al., 2018a; Ridding & Ziemann, 2010), these have included anatomical factors such as differences in skull thickness and morphology influencing the tDCS magnitude passing to the cortical area of interest (Datta et al., 2012; Miranda et al., 2013; Opitz et al., 2015). Physiological factors such as time-of-day (M. V. Sale et al., 2008; Martin V. Sale et al., 2007), hormone fluctuations with the female menstrual cycle (Inghilleri et al., 2004; Smith et al., 2002; Zoghi et al., 2015), history of past synaptic activity (Gentner et al., 2008; Iezzi et al., 2008; Rosenkranz et al., 2007) and caffeine consumption (Cerqueira et al., 2006; Hanajima et al., 2019; Orth et al., 2005). Biological factors including differences in age (Fujiyama et al., 2014; Heise et al., 2014), gender (Inghilleri et al., 2004; Zoghi et al., 2015) and specific genetic variants (Antal et al., 2010; Hwang et al., 2015; Puri et al., 2015). With inter-individual variability to tDCS still remaining, it is likely that mechanisms behind inter-individual variability are multifactorial (Li et al., 2015; Pellegrini et al., 2018a; Ridding & Ziemann, 2010). Ongoing investigation into the contribution these intrinsic factors may play in tDCS inter-individual variability is crucial.

Investigating intrinsic factors that may differ between individuals and that may serve as a predictive tool are important steps forward for developing reliable, predictable and individualised tDCS protocols for application in therapeutic and rehabilitative settings.

Genetic variations in the gene encoding brain-derived neurotrophic factor (BDNF) is one potential source of inter-individual variability previously investigated. One of the most abundant neurotrophins in the brain (Hwang et al., 2015), BDNF plays a role in activity-dependent LTP and LTD consolidation and strengthening neuronal synapses (Aicardi et al., 2004; McAllister et al., 1998; Poo, 2001). BDNF gene single nucleotide polymorphism (SNP) has been reported to influence changes in cortico-cortical excitability and CSE following NIBS. When compared to individuals with normal expression of the BDNF gene, those with the BDNF gene variant respond differently to NIBS protocols with smaller changes in CSE following tDCS (Antal et al., 2010; Frazer et al., 2016; Puri et al., 2015; Teo et al., 2014) and TBS (Antal et al., 2010; Cheeran et al., 2008; M. Lee et al., 2013). The relationship between NIBS after-effects and BDNF gene variant however is not entirely clear, with several studies reporting no association between BDNF gene expression and the after-effects of tDCS (Brunoni et al., 2013; Chhabra et al., 2016; Di Lazzaro et al., 2012; Fujiyama et al., 2014) or TBS (Li Voti et al., 2011; Mastroeni et al., 2013). These inconsistencies in research findings suggest isolated variations in BDNF gene expression may not predict inter-individual variability to NIBS.

This provides an opportunity to investigate other gene variations that may differ between individuals that more closely align with the LTP/LTD-like mechanisms of tDCS. LTP synaptic changes are mediated by the glutamate neurotransmitter and NMDA receptors (Dingledine et al., 1999; Mori et al., 2011; Nicoll & Malenka, 1999; Michael A. Nitsche et al., 2012) and the after-effects of NIBS protocols such as a-tDCS are dependent on NMDA receptor changes (Liebetanz, 2002; Michael A Nitsche et al., 2004; Michael A. Nitsche et al., 2012). Variants in the genes that encode for NMDA receptors along excitatory cortical pathways may in-part contribute to a-tDCS inter-individual variability. Additionally, with cortical excitation also regulated by gamma aminobutyric acid (GABA) inhibitory pathways (Murakami et al., 2012; Michael A. Nitsche et al., 2004, 2012), variants in the genes that encode for inhibitory GABA receptors may also contribute to a-tDCS inter-individual variability. Novel investigations into variations in genes encoding for these key regulators, and their predictive value are yet to be conducted in the tDCS literature.

This study therefore aimed to be the first of its kind to investigate the relationship between specific variants in NMDA and GABA receptors and a-tDCS inter-individual variability, and whether the selected gene variants could predict response to a-tDCS. This study will be the first of its kind in the tDCS literature to thoroughly investigate multiple NMDA and GABA receptor gene variants and their predictive value to whether an individual responds as expected or not to a-tDCS. The significance of this study will lie in its large-scale nature and its thorough investigation into multiple genetic variants and the link between inherent genetic variations and the ability to undergo cortical plasticity following a-tDCS. Conducting this study on a large sample-size will increase the power of our findings, reduce the likelihood of reporting false positive results (Minarik et al., 2016) and ensure sufficient responders and non-responders to conduct meaningful statistical analyses (Strube et al., 2016).

We hypothesised there would be an association and predictive capabilities between the selected gene variants and response to a-tDCS. We hypothesised normal expression of genes encoding for NMDA receptors would be associated with increases in CSE and a-tDCS ‘responder’ categorisation. Conversely, we hypothesised variations in genes encoding for NMDA receptors would be associated with reductions in CSE and a-tDCS ‘non-responder’ categorisation. We also hypothesised there would be an association between normal expression of genes encoding for GABA receptors and a-tDCS ‘non-responder’ categorisation and an association between GABA receptor gene variant expression and a-tDCS ‘responder’ categorisation. Lastly, we investigated the relationship between changes in intracortical inhibition (ICI) and intracortical facilitation (ICF), measures of cortico-cortical excitability, and a-tDCS response categorisation and whether they differed between a-tDCS responders and non-responders.

## Materials and methods

### Subjects

This study was approved by the human research ethics committee of Monash University according to the Declaration of Helsinki. Sixty-one healthy male volunteers aged (mean±SD) 26.82±7.62 years provided written informed consent to attend two experimental sessions. Sample-size was calculated (with 80% power) based on results of pilot (n=15). At 5% significance level, effect size of 0.45 required sample-sizes between 17-26 (Portney & Watkins, 2009). Given the subgrouping nature of this study, the number of recruited subjects was adjusted to account for identification of responders and non-responders. Previous literature report the proportion of responders range from 20-60%, with average responder rates approximately 45% (Ammann et al., 2017; Chew et al., 2015; López-Alonso et al., 2014; Puri et al., 2015, 2016; Strube et al., 2015, 2016; Tremblay et al., 2016; Wiethoff et al., 2014). To ensure the numbers of responders was between 17-26, the required sample-size was adjusted to at least fifty-eight (i.e. 26/0.45=58). Female participants were not recruited to maximise subject homogeneity and control for potential effects of fluctuating estrogen and progesterone hormones across the menstrual cycle (Inghilleri et al., 2004; Smith et al., 2002; Zoghi et al., 2015). Fifty-four subjects were right-handed and seven left-handed as measured by the Edinburgh Handedness Inventory (Oldfield, 1971) and no subject reported neurological or psychological conditions (Keel et al., 2001). All subjects were asked to refrain from drinking coffee or energy drinks at least 12-hours prior to participating (Chew et al., 2015; Fujiyama et al., 2017; Hermsen et al., 2016; Matamala et al., 2018; O’leary et al., 2015) to minimise potential confounding effects of caffeine (Cerqueira et al., 2006; Concerto et al., 2017).

### Study Design

A repeated-measures randomised cross-over design was utilised. Each subject attended two sessions in randomised order. Each session was identical except for tDCS intervention (a-tDCS or sham-tDCS). The sessions were conducted at a similar time-of-day to reduce cortisol diurnal effects (M. V. Sale et al., 2008; Martin V. Sale et al., 2007) and separated by at least one-week washing-out period to ensure no carry-over effects (Michael A. Nitsche et al., 2008).

### Electromyography Recording

Electromyography (EMG) was recorded from the dominant first dorsal interossei (FDI) via pre-gelled self-adhesive bipolar Ag/AgCl disposable surface electrodes. Skin was abraded and cleaned to minimise skin impedance (Gilmore & Meyers, 1983). Electrodes were placed over the FDI with 2cm inter-electrode distance and reference electrode placed over ulna styloid process (Kendell et al., 2010). EMG signals were filtered, amplified (10-500Hz x 1000) and sampled at 1000Hz. EMG data were collected via a laboratory analogue-digital interface (LabChart^TM^ & Powerlab, ADInstruments, Australia).

### Establishment of Intra-Rater Reliability for assessment of corticospinal excitability

Single-assessor intra-rater reliability for the assessment of corticospinal excitability via the TMS device has been previously established (Pellegrini et al., 2018c). CSE as measured by TMS-evoked MEPs were recorded over a range of TMS test intensities; 105%, 120%, 135%, 150%, 165% of resting motor threshold (RMT). Ranging from 0.660-0.968, significant (p<0.05) inter-class correlations were reported both within-sessions and between-sessions for TMS test intensities 120-165% of RMT (Pellegrini et al., 2018c).

### Transcranial Magnetic Stimulation Procedure

Single and paired-pulse TMS paradigms were delivered to the dominant primary motor cortex (M1) by a 70mm figure-of-eight TMS coil (Magstim Limited Company, UK). The coil was held over M1 oriented 45° to the midline and tangential to the scalp for posterior-anterior current flow (Rossini & Rossi, 1998) and manually repositioned to determine the cortical area to elicit the greatest FDI motor response. This hotspot was marked on the scalp to ensure consistent placement throughout the session.

RMT was defined as the percentage of maximal stimulator output (MSO) required for MEP peak-to-peak amplitude greater than 50µV in 5/10 consecutive stimuli (Devanne et al., 2006). Test intensity was defined as the percentage of TMS MSO required to elicit an MEP peak-to-peak amplitude of ∼1mV. Starting at 50% MSO, the intensity was adjusted in 1-2% intervals until the above RMT and test intensity criteria were met (Rothwell et al., 1999). These intensities were calculated pre and post tDCS intervention for each session with RMT used to calculate conditioning stimuli for paired-pulse TMS paradigms.

### Outcome Measures

As an index of CSE, twenty-five single-pulse MEPs with 6sec inter-trial interval (ITI) at the test intensity were recorded. Paired-pulse TMS paradigms were used for the assessment of cortico-cortical excitability. A conditioning pulse (80% RMT) followed by a pulse at the test intensity were separated by 3msec inter-stimulus interval (ISI) for short-interval intracortical inhibition (SICI) and 10msec for ICF (Di Pino et al., 2014; Kujirai et al., 1993). Fifty paired-pulse MEPs, 25 with 3msec and 10msec ISI, were delivered in randomised order separated by 6sec ITI. Considered an index of M1 ICI and ICF, mean values were calculated for SICI and ICF and expressed as percentages of single-pulse mean MEP (Di Pino et al., 2014; Kujirai et al., 1993).

### Transcranial Direct Current Stimulation

TDCS was delivered with current intensity and duration of 1mA and 10-minutes with 30-second fade-in/fade-out periods (NeuroConn DC-stimulator, Germany). Delivered via two rectangular saline-soaked sponge electrodes fixed to the scalp by velcro straps, the active electrode (4×6cm, 24cm^2^) was centred over the dominant M1 with return electrode (5×7cm, 35cm^2^) over the contralateral supraorbital area. Current density was 0.0417mA/cm^2^ under the active electrode and 0.0286mA/cm^2^ under the return electrode, with the size and current density differential between active and return electrodes focussing electric current under the active electrode and away from the return electrode (M. A. Nitsche et al., 2007b). For sham-tDCS, the electrode over the contralateral supraorbital area was the active electrode with current increasing from 0-1mA for a 30-second fade-in period then reducing to 0mA for the remaining 9.5-minutes. Participants were blinded to the nature of each intervention (a-tDCS and sham-tDCS). Blinding integrity was assessed by asking participants the nature of the intervention at the conclusion of each session.

Although tDCS is considered a safe intervention with no serious adverse effects (Michael A. Nitsche et al., 2008), the tolerability and side-effects to tDCS were assessed both during and after the application of tDCS. The presence of any sensation of itching, tingling or discomfort were monitored during the application of tDCS. Participants were asked to rate the presence of sensations on a scale of 1-10 at the beginning and middle of tDCS application. The presence of headache or other sensory complaints following tDCS were also assessed on a scale of 1-10.

### Genotyping Procedure

Saliva samples were obtained via Oragene-DNA self-collection kits (DNAgenotek, Ontario, Canada). Ten genetic variations were chosen where a single nucleotide was substituted for another in a particular stretch of DNA. Ten SNPs were selected based on their involvement in synaptic transmission. Based on previous literature, SNPs for the BDNF gene (Antal et al., 2010; Cheeran et al., 2008; Di Lazzaro et al., 2012; Frazer et al., 2016; Puri et al., 2015) and glutamate NMDA receptor genes GRIN1 (L.-C. Lee et al., 2016; Mori et al., 2011; Rossi et al., 2013) and GRIN2B (Mori et al., 2011; Narita et al., 2018) were selected for their involvement in excitatory glutamatergic cortical pathways while GABA receptor genes GABRA1, GABRA2 and GABRA3 (Hung et al., 2013) were selected for their involvement in inhibitory GABAergic cortical pathways (table 1). Genotyping was performed by the Australian Genome Research Facility (AGRF, St Lucia, Queensland). For each gene, subjects were classified into either having ‘normal expression’ (i.e. common homozygote) or ‘variant expression’ (i.e. heterozygous or homozygous for the substituted nucleotide). To avoid assessor bias, genotype analysis was not conducted until all other data collection was completed.

**Table 1.**
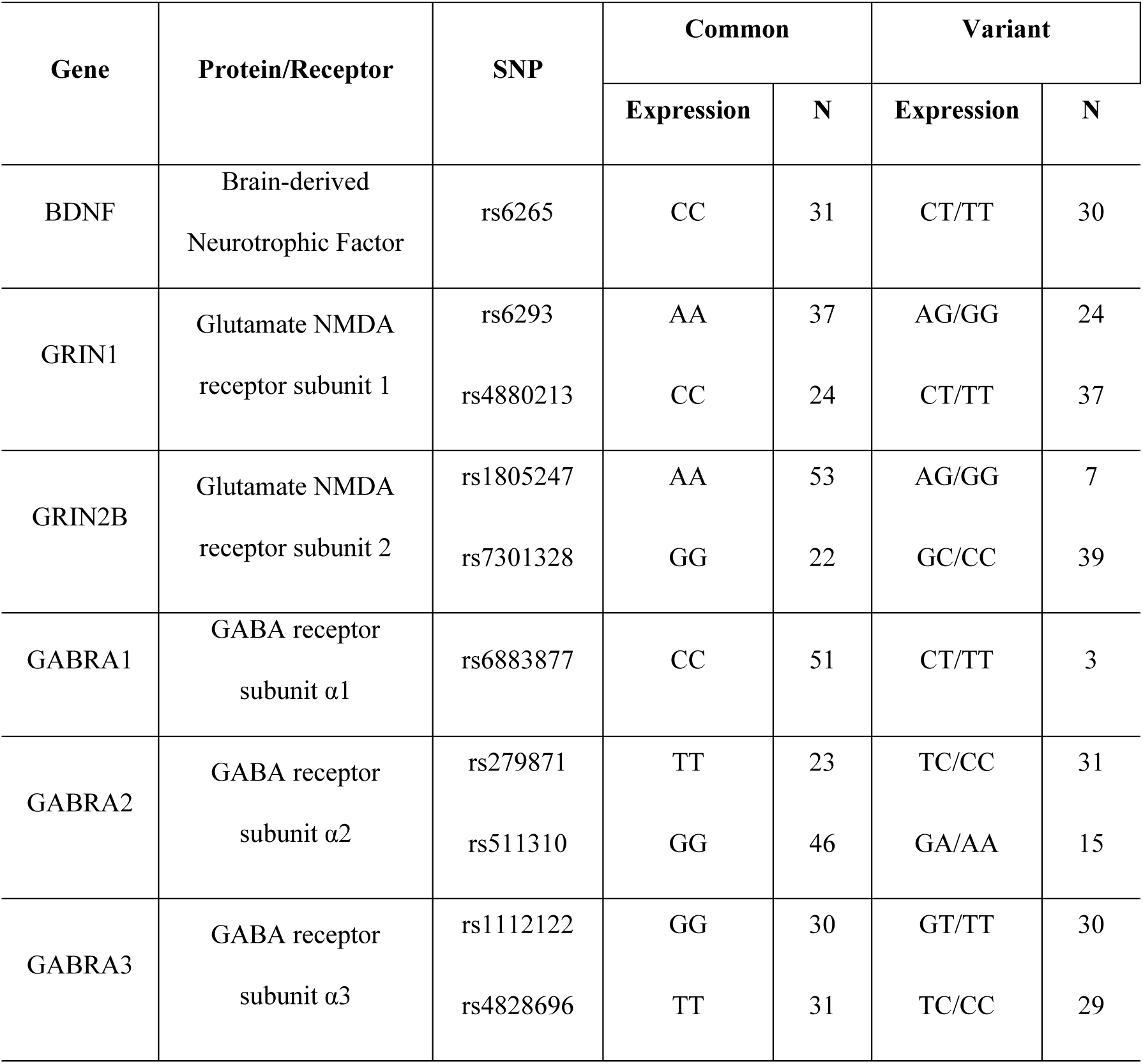
Selected genes and SNPs. Summary of selected genes and SNPs with normal expression (common homozygous) and variant expression (heterozygous and rare homozygous) with accompanying sample-sizes.

### Experimental Procedure

The sessions (a-tDCS, sham-tDCS) were conducted in randomised order. Subjects sat relaxed in an adjustable chair, their hand at rest. RMT and test intensities were determined followed by, baseline measures data collection. TDCS was then administered, immediately followed by single-pulse MEP data collection at 0-minutes post-tDCS. RMT and test intensity were then re-established for SICI and ICF data collection at 10-minutes post-tDCS. Single-pulse MEPs were again recorded at 30-minutes post-tDCS, while SICI and ICF data were re-recorded at 40-minutes post-tDCS to investigate whether measures measured had return to baseline values (figure 1) (A. Bastani & Jaberzadeh, 2013; Andisheh Bastani & Jaberzadeh, 2013). Saliva samples were collected at the conclusion of the second session.

**Figure 1.**
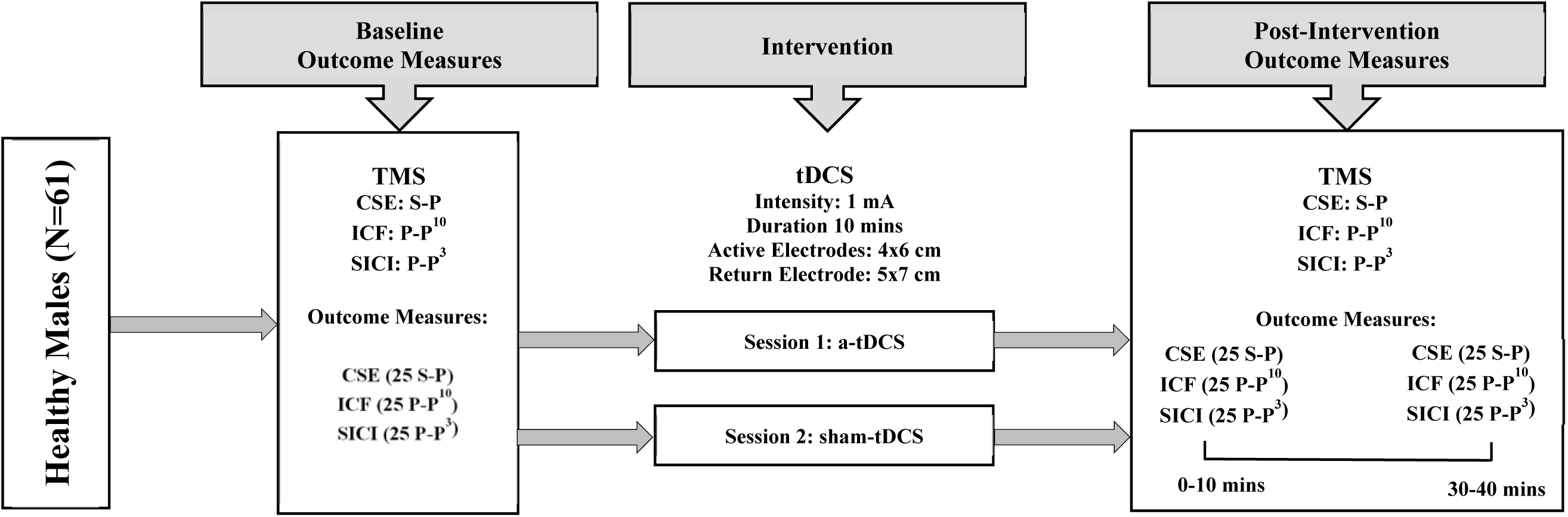
Experimental design. 25 single-pulse and 50 paired-pulse TMS MEPs delivered to M1 hotspot for FDI. Both a-tDCS and sham-tDCS delivered on separate testing sessions before re-assessing outcome measures at 0-10 minutes and 30-40 minutes post-tDCS.

### Statistical Analyses

Statistical analyses were conducted using SPSS (Version 25.0, IL, USA).

### Group level analysis

All outcome measures (MEP, SICI, ICF) were assessed for each intervention (a-tDCS, sham-tDCS) and time-point (baseline, 0-minutes, 30-minutes). Shapiro-Wilk test assessed data normality for each outcome measure. Non-parametric statistical tests were used if data violated normality.

Between-session reliability via two-way repeated measures analysis of variance (RM-ANOVA) assessed cumulative effects of changes in each outcome measure (MEP, SICI, ICF) in repeated measures study designs (Alonzo et al., 2012; Gálvez et al., 2013; Michael A. Nitsche et al., 2008). Baseline values for each session assessed the effect of intervention on within-subject reliability to ensure no carry-over effects between sessions (Portney & Watkins, 2009). The main effects of time, intervention and their interaction were then investigated. For CSE, as measured by single-pulse MEP, two-way repeated-measures analysis of variance (RM-ANOVA) investigated overall effects of time, intervention and their interaction on MEP. Time (three levels) and intervention (two levels) were the independent factors. Mauchly’s sphericity test examined equality of variances, with Greenhouse-Geisser’s correction utilised when variances were unequal and sphericity was not assumed. Post-hoc comparisons using the Bonferroni correction and one-way RM-ANOVA were utilised where indicated to test whether post a-tDCS MEP were significantly different from baseline and significantly different from sham-tDCS at each time-point.

The non-parametric Friedman’s two-way ANOVA was used to investigate the overall effects of time and intervention on SICI and ICF data which violated normality. The non-parametric Wilcoxon matched-paired test was then used post-hoc where indicated to investigate the SICI and ICF values at post-tDCS time-points were significantly different from baseline values following both a-tDCS and sham-tDCS. All tests were performed at 5% significance level (α=0.05).

### Subgroup level analysis

Two techniques were used to categorise subjects into subgroups. Firstly, subjects were categorised based on whether their post-tDCS responses exceeded a pre-determined threshold. MEPs at 0-minutes post-tDCS were normalised to baseline and compared to the pre-determined threshold of the SD of sham-tDCS baseline values as previously used (Ammann et al., 2017). This selected threshold accounted for the natural variability in response within the current cohort of subjects and accounted for inherent variability in measuring CSE via TMS-evoked MEPs. Given MEPs were collected at only two post-tDCS time-points (0-minutes, 30-minutes), taking a grand average of all post-tDCS time-points as previously conducted (Ammann et al., 2017; Chew et al., 2015; López-Alonso et al., 2014; Puri et al., 2015, 2016; Tremblay et al., 2016; Wiethoff et al., 2014) would not have provided a true indication of how each subject immediately responded to a-tDCS as after-effects have been reported to return to baseline levels at 30-minutes post-tDCS (A. Bastani & Jaberzadeh, 2013; Andisheh Bastani & Jaberzadeh, 2013). Baseline sham-tDCS SD was calculated at 0.324, therefore based on their normalised MEPs, subjects were classified as either ‘responders’ (post a-tDCS nMEP>1.324) or non-responders (post a-tDCS nMEP<1.324).

SPSS TwoStep cluster analysis was the second subgrouping technique utilised. This technique facilitated subgrouping based on existing trends or patterns in the data (Hair et al., 1998) and its use in NIBS literature is reviewed elsewhere (Pellegrini et al., 2018b). Cluster predictors were normalised MEP at 0-minutes post a-tDCS.

For each of the two subgroups identified by the subgrouping techniques (i.e. responders/non-responders and cluster 1/cluster 2), one-way ANOVA were conducted to investigate significant differences between subgroups in MEP amplitude at each time-point. Subgroups were then analysed independently to investigate the effect of time and intervention on SICI and ICF data via non-parametric Friedman’s and Wilcoxon matched-paired tests. This investigated whether post-tDCS changes in ICI or ICF differed between a-tDCS responders/cluster 1 compared to a-tDCS non-responders/cluster 2. Non-parametric Mann-Whitney U test then determined significant differences between subgroups at any post-tDCS time-point. Lastly, similarities between subgrouping techniques were investigated via chi-squared tests to assess for significant associations between the two different subgrouping techniques.

### Genotype analysis

Genotype data was analysed against subgroup categories (i.e responders/non-responders and cluster 1/cluster 2) to investigate associations between gene expression and a-tDCS response and whether gene expression is a predictor of a-tDCS response. Chi-squared tests analysed each gene individually against subgroup category and cluster membership testing for associations or dependency between each gene expression and a-tDCS response. Univariate binary logistical regression analysis was then conducted for each gene variant independently to quantify the association (i.e. Odds ratio) between subgroup category and gene expression to assess the predictive value of each gene variant for a-tDCS response. Subgroup category was the dependent variable while each gene was the independent variable. Multivariate binary logistical regression analysis was then conducted as above to investigate associations and predictive value of each gene variant for a-tDCS response when controlling and accounting for all other gene variants.

If significant associations were revealed between a particular gene variant and a-tDCS response, individuals were subgrouped as either ‘normal expression’ or ‘variant expression’ and analysed independently. Two-way and one-way RM-ANOVA on MEP data were conducted to determine the post-tDCS time-point where MEP significantly differed from baseline and at which post-tDCS time-point MEP values between ‘normal expression’ and ‘variant expression’ subgroups significantly differed. Non-parametric Friedman’s and Wilcoxon matched-paired and Mann-Whitney U tests were also conducted independently in genetic ‘normal expression’ and ‘variant expression’ subgroups to investigate differences between genetic subgroups in SICI and ICF data.

## Results

All sixty-one participants attempted each session with mean (±SD) 13.58±15.91 days between each session. Mean RMT and test intensity were 36% (36.29±7.69%) and 46% (45.78±10.10%) of the TMS device MSO respectively.

### Tests for Normality of data

Shapiro-Wilk tests revealed single-pulse MEP peak-to-peak amplitude data were normally distributed for both tDCS interventions and all time-points (p>0.05). SICI and ICF data violated normality tests (p<0.05).

### Carry-Over effects

RM-ANOVA revealed no significant differences in baseline values between a-tDCS and sham-tDCS for MEP (p=0.200). Friedman’s test revealed no significant differences in baseline values between a-tDCS and sham-tDCS for SICI (p=0.055) and ICF (p=0.249). These results indicate no carry-over effects between sessions.

### Group level analysis

#### Corticospinal Excitability

Two-way RM-ANOVA showed significant main effects of time (p<0.01), intervention (p<0.01) and time*intervention interaction (p<0.01). Post-hoc analysis and one-way RM-ANOVA revealed when compared to baseline, MEP amplitudes significantly increased at 0-minutes following a-tDCS (p<0.05), significantly greater than MEP amplitudes following sham-tDCS at the same time-point (p<0.01). MEP amplitude then reduced towards baseline at 30-minutes following a-tDCS (p>0.05). No significant changes were revealed for sham-tDCS between time-points (p>0.05) (figure 2a, 2b and table 2).

**Figure 2.**
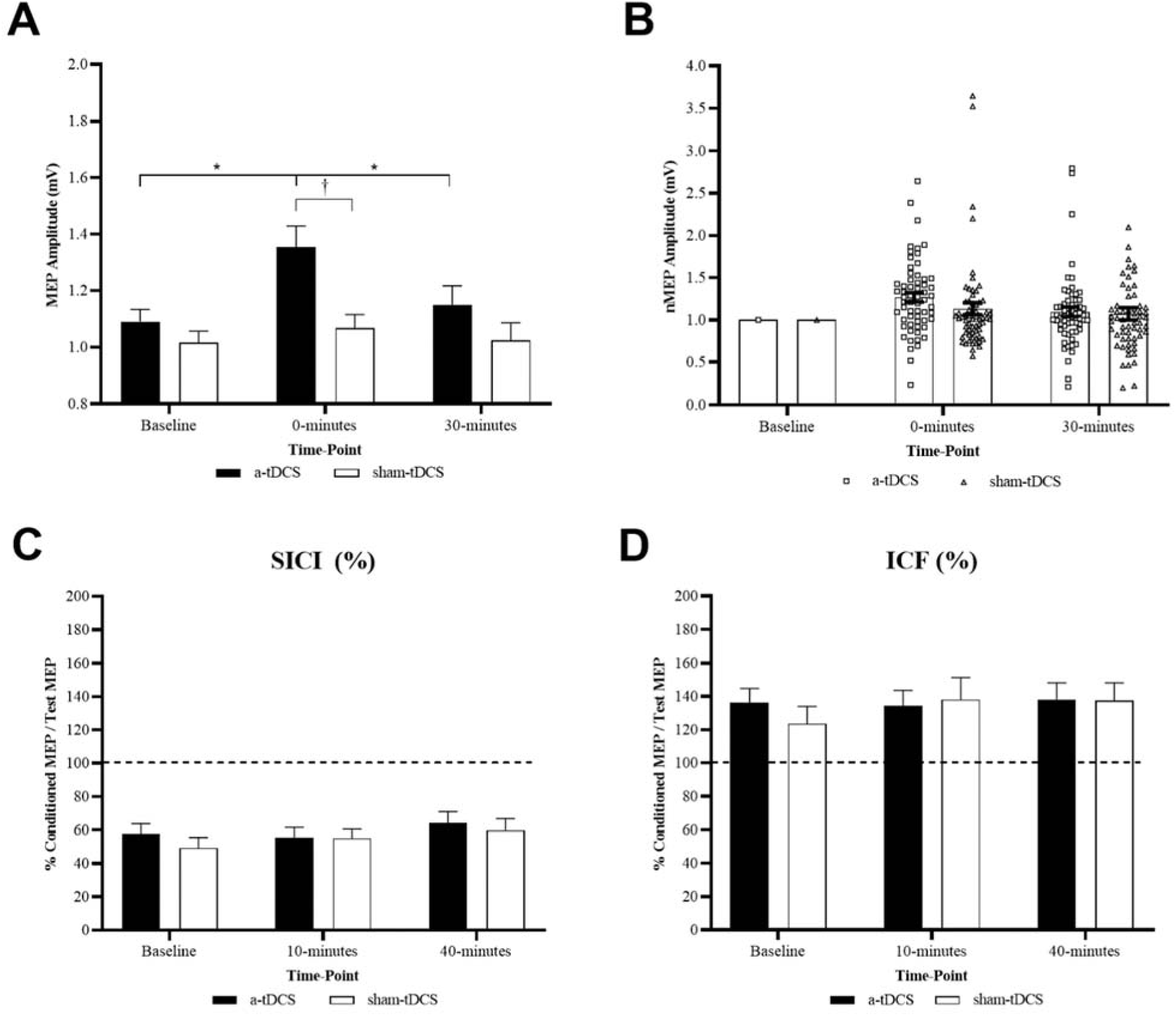
Group level a-tDCS versus sham-tDCS for each time-point. (a) Mean single-pulse MEP amplitudes. (b) Single-pulse normalised MEP amplitude with individual data points. (c) ICI as measured by SICI values. (d) ICF. Error bars denote one SEM. *denote significant differences between time-points (p<0.05). ^†^denote significant differences between interventions at a particular time-point (p<0.01).

**Table 2.** Group level corticospinal excitability and cortico-cortical excitability. Mean(±SD) MEP peak-to-peak amplitude (mV), SICI (%) and ICF (%) data for each intervention and time-point. Post-tDCS time-point 1 is 0-minutes for MEP and 10-minutes for SICI and ICF. Post-tDCS time-point 2 is 30-minutes for MEP and 40-minutes for SICI and ICF. *denote significant between post-tDCS time-points and baseline (p<0.05). ^†^denote significant differences between interventions at a particular time-point (p<0.05).

| **Intervention** | **Outcome Measure** | **Baseline** | **Post-tDCS time-point 1** | **Post-tDCS time-point 2** |
| --- | --- | --- | --- | --- |
| **a-tDCS** | **MEP (mV)** | 1.090±0.337 | 1.354±0.587 *† | 1.150±0.522 |
| | **SICI (%)** | 57.72±47.37% | 55.49±47.92% | 64.09±53.31% |
| | **ICF (%)** | 136.22±67.23% | 134.27±73.80% | 138.20±77.62% |
| **sham-tDCS** | **MEP (mV)** | 1.015±0.324 | 1.066±0.377 † | 1.024±0.487 |
| | **SICI (%)** | 49.28±49.78% | 55.22±41.02% | 59.67±55.50% |
| | **ICF (%)** | 123.71±81.01% | 138.13±103.22% | 137.66±81.51% |

#### Cortico-cortical Excitability

Friedman’s test showed no significant effects of time or intervention for SICI and ICF (p>0.05) (figures 2c, 2d and table 2).

### Subgroup level analysis

#### Subgrouping based on a pre-determined threshold

Subgrouping based on each subjects normalised responses at 0-minutes post a-tDCS with a 32.4% threshold revealed two different subgroups. Following a-tDCS, 43% (n=26) of subjects were categorised as ‘responders’ (nMEP>1.324 at 0-minutes post a-tDCS) while 57% (n=35) were categorised as ‘non-responders’ (nMEP<1.324 at 0-minutes post a-tDCS) (figures 3c).

**Figure 3.**
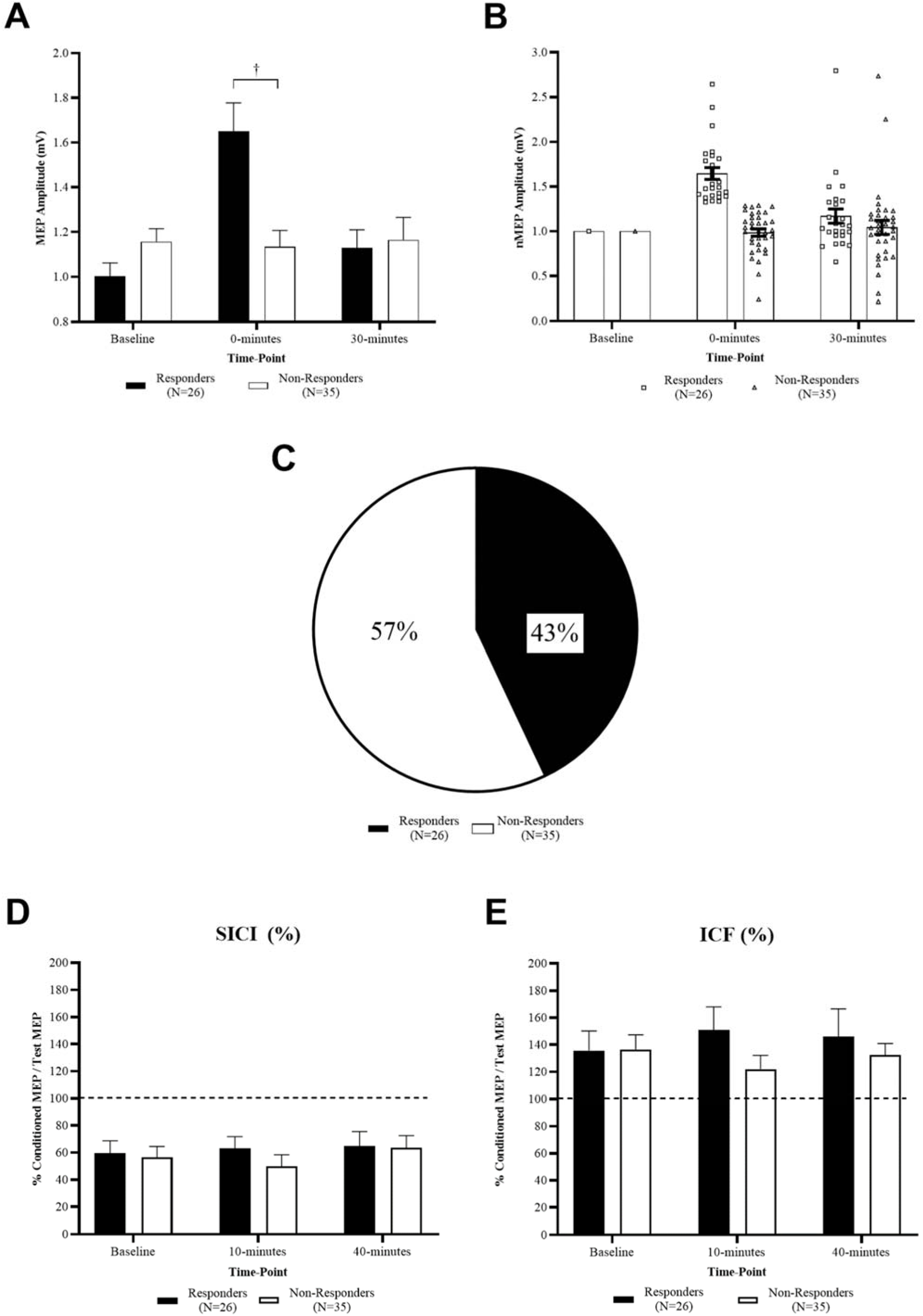
Subgrouping based on pre-determined threshold. Responders versus Non-Responders before and after a-tDCS at each time-point. (a) Single-pulse MEP amplitudes. (b) Single-pulse normalised MEP amplitude with individual data points. (c) Percentage breakdown of responders and non-responders. (d) ICI as measured by SICI. (e) ICF. Error bars denote one SEM. ^†^denote significant differences between responder and non-responder subgroups at a particular time-point (p<0.05).

#### Corticospinal Excitability

Baseline MEP amplitudes were not significantly different between responder and non-responder subgroups for both a-tDCS and sham-tDCS (p>0.05) or between a-tDCS and sham-tDCS within responder and non-responder subgroups (p>0.05). One-way ANOVA revealed that at 0-minutes following a-tDCS, MEP amplitude was significantly greater in responders compared to non-responders (p<0.01). No significant differences were revealed between subgroups at 30-minutes following a-tDCS (p>0.05) or following sham-tDCS (p>0.05) (figures 3a-b and table 3).

**Table 3.** Corticospinal excitability for responders/non-responders, clusters 1/cluster 2 and GABRA3 gene rs4828696 SNP normal/variant expression subgroups. Mean(±SD) MEP peak-to-peak amplitude (mV) for both interventions and time-point. *denote significant difference within a subgroup between a post-tDCS time-point and baseline (p<0.05). ^†^denote significant difference between subgroups at a particular time-point (p<0.05). ^#^denote significant difference between tDCS interventions for a particular subgroup (p<0.05).

| Intervention | Category | Baseline | 0-minutes | 30-minutes |
| --- | --- | --- | --- | --- |
| a-tDCS | Responders | 1.001±0.308 | 1.651±0.641 *†# | 1.130±0.409 |
| | Non-Responders | 1.157±0.346 | 1.134±0.434 † | 1.165±0.598 |
| | Cluster 1 | 0.984±0.321 | 1.667±0.678 *†# | 1.122±0.434 |
| | Cluster 2 | 1.115±0.333 | 1.163±0.431 †# | 1.167±0.574 |
| | GABRA3 rs4828696<br>'Variant Expression' | 1.019±0.354 | 1.348±0.671 * | 1.041±0.367 |
| | GABRA3 rs4828696<br>'Normal Expression' | 1.164±0.312 | 1.374±0.512 * | 1.268±0.620 |
| sham-tDCS | Responders | 0.943±0.302 | 0.970±0.338 # | 0.954±0.459 |
| | Non-Responders | 1.068±0.333 | 1.138±0.393 | 1.076±0.507 |
| | Cluster 1 | 0.968±0.271 | 0.989±0.341 # | 0.983±0.469 |
| | Cluster 2 | 1.043±0.352 | 1.113±0.394 | 1.048±0.502 |
| | GABRA3 rs4828696<br>'Variant Expression' | 0.984±0.346 | 1.067±0.387 | 0.992±0.457 |
| | GABRA3 rs4828696<br>'Normal Expression' | 1.045±0.310 | 1.068±0.380 # | 1.059±0.525 |

#### Cortico-cortical Excitability

For SICI, there were no significant differences at baseline between responders and non-responders for a-tDCS and sham-tDCS (p>0.05). For responders, no significant differences were reported at baseline between a-tDCS and sham-tDCS (p>0.05). For non-responders, baseline values differed between a-tDCS and sham-tDCS (p<0.05). Friedman’s and Wilcoxon matched-paired test revealed no significant changes in SICI for both responder and non-responder subgroups following both a-tDCS and sham-tDCS (p>0.05) or between a-tDCS and sham-tDCS interventions (p>0.05). For a-tDCS, Mann-Whitney U tests revealed no significant differences between subgroups at either post a-tDCS time-points (p>0.05). For sham-tDCS, significant differences between subgroups were revealed at 10-minutes post sham-tDCS (p<0.05) but not at 40-minutes post sham-tDCS (p>0.05) (figure 3e and table 4).

**Table 4.** Cortico-cortical excitability for responders and non-responders subgroups. Mean(±SD) SICI and ICF data for each intervention and time-point.

| Outcome Measure | Subgroup | SICI (%) |  |  | ICF (%) |  |  |
| --- | --- | --- | --- | --- | --- | --- | --- |
|  |  | Baseline | 10-minutes | 40-minutes | Baseline | 10-minutes | 40-minutes |
| a-tDCS | Responders | 59.42±47.11% | 63.24±43.37% | 64.90±54.33% | 135.72±73.42% | 150.78±87.90% | 146.10±104.20% |
| | Non-Responders | 56.46±48.20% | 49.74±50.89% | 63.48±53.33% | 136.59±63.43% | 122.01±59.72% | 132.33±50.65% |
| sham-tDCS | Responders | 49.60±35.90% | 68.77±41.20% | 67.99±68.95% | 118.74±53.23% | 159.78±138.46% | 148.26±98.20% |
| | Non-Responders | 49.05±58.52% | 45.16±38.43% | 53.50±42.99% | 127.40±97.29% | 122.04±63.88% | 129.79±66.95% |

For ICF data, there were no significant differences in baseline values between responders and non-responders for a-tDCS and sham-tDCS (p>0.05). No significant differences were reported at baseline between a-tDCS and sham-tDCS for both responders and non-responders (p>0.05).

Friedman’s and Wilcoxon matched-paired test revealed no significant changes in ICF for both responder and non-responder subgroups following both a-tDCS and sham-tDCS (p>0.05) or between a-tDCS and sham-tDCS interventions (p>0.05). Mann-Whitney U tests revealed no significant differences between subgroups at any of the post-tDCS time-points for both a-tDCS and sham-tDCS (p>0.05) (figure 3f and table 4).

### Two-Step Cluster Analysis

Two-step cluster analysis revealed two distinct clusters of data. Cluster 1 encompassed 38% (n=23) of subjects that responded to a-tDCS with increases in MEP amplitude. Cluster 2 included the remaining 62% (n=38) of subjects that responded to a-tDCS with no increase or reductions in MEP amplitude (figure 4c).

**Figure 4.**
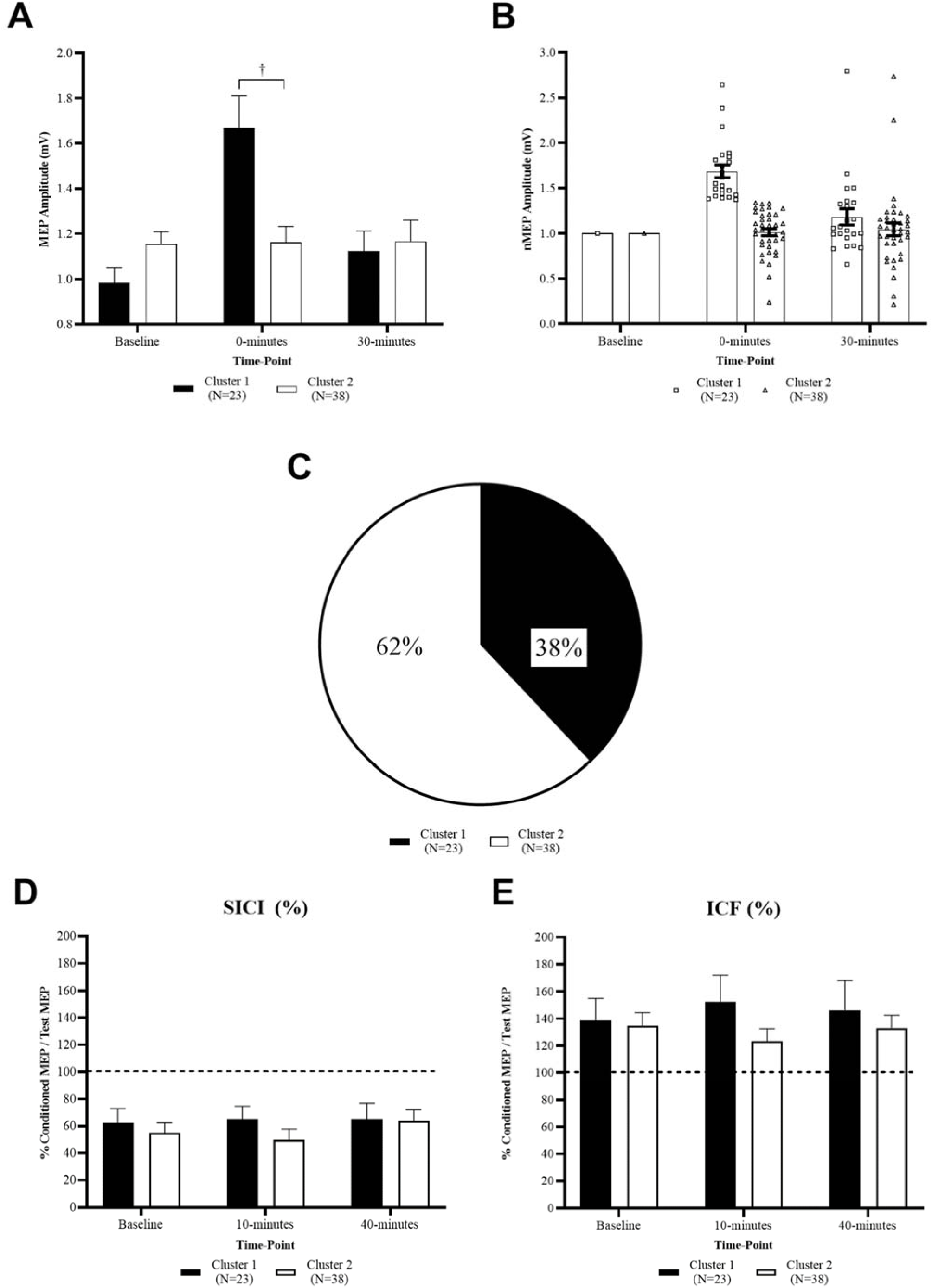
Subgrouping based on cluster analysis. Cluster 1 versus Cluster 2 before and after a-tDCS at each time-point. (a) Single-pulse MEP amplitudes. (b) Single-pulse normalised MEP amplitude with individual data points. (c) Percentage breakdown of cluster 1 and cluster 2. (d) ICI as measured by SICI. (e) ICF. Error bars denote one SEM. ^†^denote significant difference between cluster 1 and cluster 2 subgroups at a particular time-point (p<0.05).

#### Corticospinal Excitability

Baseline MEP amplitudes were not significantly different between cluster 1 and cluster 2 subgroups for either a-tDCS or sham-tDCS (p>0.05) or between a-tDCS and sham-tDCS within both cluster subgroups (p>0.05). One-way ANOVA revealed at 0-minutes following a-tDCS, MEP amplitude was significantly greater in cluster 1 compared to cluster 2 (p<0.01). No significant differences were revealed between cluster subgroups at 30-minutes following a-tDCS (p>0.05) or at all sham-tDCS time-points (p>0.05) (figures 4a-b and table 4).

#### Cortico-cortical Excitability

For SICI, there were no significant differences at baseline between cluster 1 and cluster 2 subgroups for a-tDCS and sham-tDCS (p>0.05). For cluster 1, significant differences were reported at baseline between a-tDCS and sham-tDCS (p<0.05). For cluster 2, baseline values were not significantly different between a-tDCS and sham-tDCS (p>0.05). Friedman’s and Wilcoxon matched-paired test revealed no significant changes in SICI for both cluster 1 and cluster 2 subgroups following both a-tDCS and sham-tDCS (p>0.05) or between a-tDCS and sham-tDCS interventions (p>0.05). For a-tDCS, Mann-Whitney U tests revealed no significant differences between cluster subgroups at either post a-tDCS time-points (p>0.05). For sham-tDCS, significant differences between cluster subgroups were revealed at 10-minutes post sham-tDCS (p<0.05) but not at 40-minutes post sham-tDCS (p>0.05) (figure 4e and table 5).

**Table 5.** Cortico-cortical excitability for cluster 1 and cluster 2 subgroups. Mean(±SD) SICI and ICF data for each intervention and time-point.

| Outcome Measure | Subgroup | SICI (%) |  |  | ICF (%) |  |  |
| --- | --- | --- | --- | --- | --- | --- | --- |
|  |  | Baseline | 10-minutes | 40-minutes | Baseline | 10-minutes | 40-minutes |
| a-tDCS | Cluster 1 | 62.50±49.27% | 64.87±45.94% | 65.10±55.89% | 138.88±77.63% | 152.52±92.95% | 146.44±103.94% |
| | Cluster 2 | 54.83±46.61% | 49.82±48.80% | 63.47±52.45% | 134.61±61.15% | 123.23±57.99% | 133.20±57.27% |
| sham-tDCS | Cluster 1 | 47.54±30.56% | 68.13±42.48% | 69.84±71.83% | 115.29±49.50% | 161.15±147.23% | 152.17±103.08% |
| | Cluster 2 | 50.34±58.82% | 47.41±38.59% | 53.52±42.72% | 128.81±95.47% | 124.19±62.12% | 128.88±65.15% |

For ICF, there were no significant differences in baseline values between cluster 1 and cluster 2 subgroups for a-tDCS and sham-tDCS (p>0.05). No significant differences were reported at baseline between a-tDCS and sham-tDCS for both clusters 1 and 2 (p>0.05). Friedman’s and Wilcoxon matched-paired test revealed no significant changes in ICF for both cluster 1 and cluster 2 subgroups following both a-tDCS and sham-tDCS (p>0.05) or between a-tDCS and sham-tDCS interventions (p>0.05). Mann-Whitney U tests revealed no significant differences between subgroups at any of the post-tDCS time-points for both a-tDCS and sham-tDCS (p>0.05) (figure 4f and table 5).

### Associations between subgrouping techniques

Twenty-three (n=23) subjects were classified as both a responder based on a pre-determined threshold and cluster 1 membership via two-step cluster analysis. Thirty-five (n=35) subjects were classified as both a non-responder and cluster 2 membership. Fisher’s exact test revealed a significant association between subgrouping based on a pre-determined threshold and two-step cluster analysis classification techniques (p<0.01).

### Genotype analysis

#### Association/dependency between gene expression and subgroup categorisation

Genotype analysis revealed seven of the ten genes had sufficient sample-sizes in both ‘normal expression’ and ‘variant expression’ subgroups to carry-out analyses (table 1). GRIN2B rs1805247, GABRA1 rs6883877 and GABRA2 rs511310 had disproportionate sample-sizes in each of the dichotomous subgroups and were therefore excluded from analysis (table 1). Chi-squared analysis revealed no significant association between subgroup category and BDNF rs6265, GRIN1 rs6293 and rs4880213, GRIN2B rs7301328 and GABRA2 rs279871 (p>0.05). Significant associations were revealed between GABRA3 rs1112122 (p=0.021) and rs4828696 (p=0.036) and subgrouping category based on pre-determined threshold, indicating dependency between the expression of GABRA3 gene SNPs rs1112122 and rs4828696 and responder/non-responder subgroup categorisation. No significant associations were reported between each gene expression and cluster 1 and cluster 2 subgroups.

#### Gene expression as predictor of response to a-tDCS

Univariate logistic regression analysis and odds ratio (OR) calculation revealed subjects with GABRA3 gene SNP rs1112122 ‘variant expression’ were three times (OR=3.051) more likely to be categorised as an a-tDCS responder based on the pre-determined threshold when compared to subjects with ‘normal expression’ of GABRA3 gene SNP rs1112122 (p=0.040). Additionally, subjects with GABRA3 gene SNP rs4828696 ‘variant expression’ were 3.5 times (OR=3.463) more likely to be categorised as an a-tDCS responder when compared to subjects with ‘normal expression’ of GABRA3 gene SNP rs4828696 (p=0.023), indicating when analysed independently, the two GABRA3 gene SNPs had significant predictive value for response to a-tDCS.

Multivariate logistic regression analysis and OR calculation revealed when all genes were controlled and accounted for, no significant relationships were reported between gene expression of the selected SNPs and subgroup category based on pre-determined threshold (p>0.05). It is worth noting however that subjects with GABRA3 gene SNP rs4828696 ‘variant expression’ were five times (OR=5.341) more likely to be categorised as an a-tDCS responder based on pre-determined threshold when compared to subjects with ‘normal expression’, a result that was approaching significance (p=0.073). Therefore when each gene variant is controlled and accounted for, no significant associations indicate no predictive value of the selected gene SNPs for subgroup response category.

#### GABRA3 SNP rs4828696 gene expression and cortico-cortical and corticospinal excitability

One-way RM-ANOVA revealed no significant effect of time for MEP amplitude following a-tDCS in GABRA3 gene SNP rs4828696 ‘normal expression’ subjects. For ‘variant expression’ however, significant effects of time were revealed for MEP amplitude following a-tDCS with significant increases at 0-minutes when compared to baseline (p<0.01) (figures 5a-b and table 6). These changes were not significantly different when compared to sham-tDCS at the same time-points (figure 5c). No significant effects of time or intervention were reported for SICI and ICF following both a-tDCS and sham-tDCS in either GABRA3 gene SNP rs4828696 ‘normal expression’ or ‘variant expression’ subgroups (figures 5d-e).

**Figure 5.**
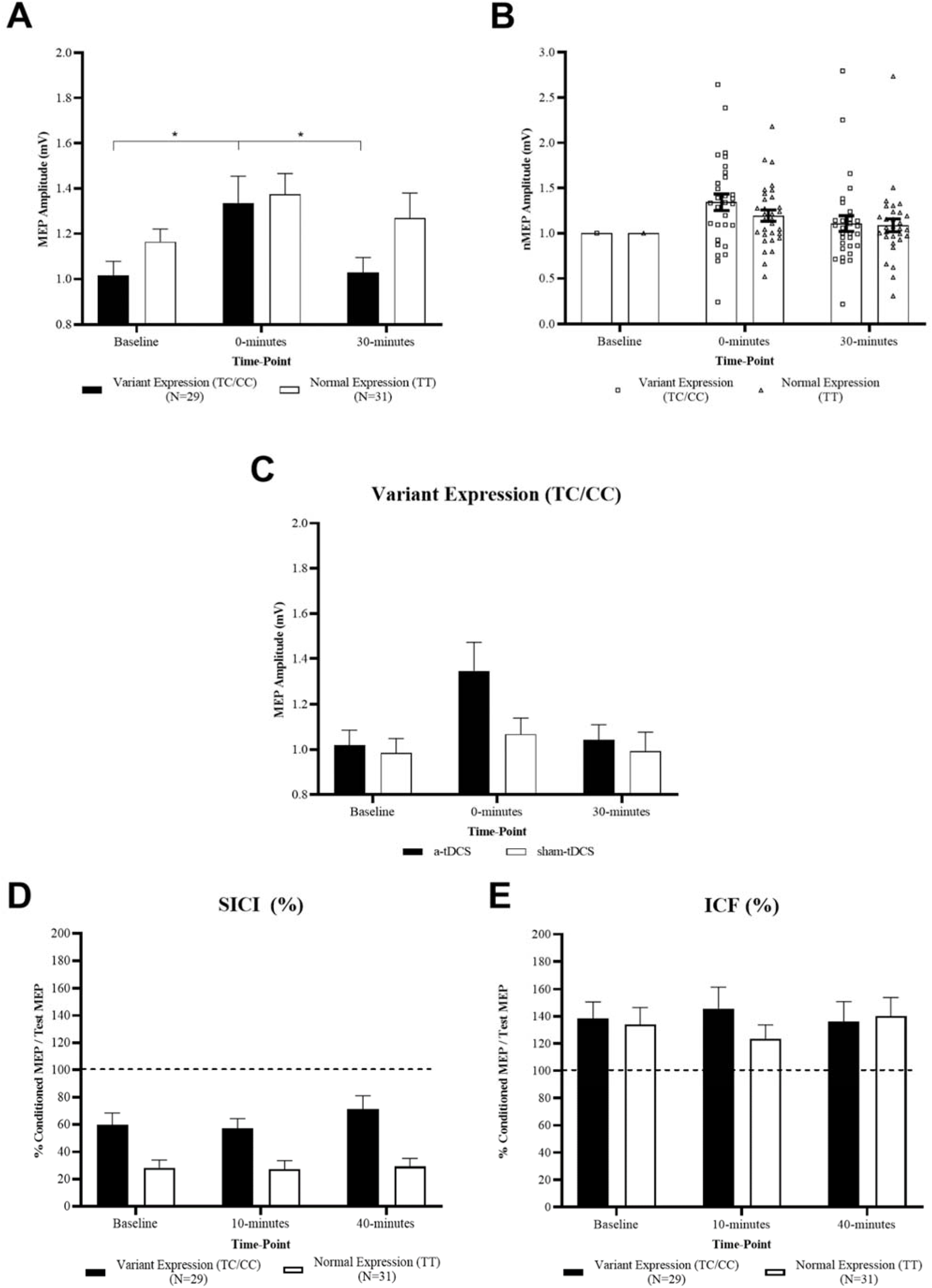
Subgrouping based on GABRA3 gene SNP rs4828696 expression. ‘Variant expression’ versus ‘Normal expression’ at each time-point. (a) Single-pulse MEP amplitudes before and after a-tDCS at each time-point. (b) Single-pulse normalised MEP amplitude with individual data points following a-tDCS. (c) Single-pulse MEP amplitude following a-tDCS and sham-tDCS in ‘Variant expression’ subgroup. (d) ICI as measured by SICI. (e) ICF. Error bars denote one SEM. *denote significant difference between time-points within a particular gene expression subgroup (p<0.05).

**Table 6.** Cortico-cortical excitability for GABRA3 gene SNP rs4828696 ‘normal expression’ and ‘variant expression’ subgroups. Mean(±SD) SICI and ICF data for each intervention and time-point.

| Outcome Measure | Subgroup | SICI (%) |  |  | ICF (%) |  |  |
| --- | --- | --- | --- | --- | --- | --- | --- |
|  |  | Baseline | 10-minutes | 40-minutes | Baseline | 10-minutes | 40-minutes |
| a-tDCS | GABRA3 rs4828696<br>‘Variant Expression’ | 57.67±45.61% | 56.18±37.14% | 67.79±52.08% | 137.65±68.02% | 147.18±87.59% | 135.09±81.73% |
| | GABRA3 rs4828696<br>‘Normal Expression’ | 55.58±48.86% | 53.65±57.06% | 57.32±52.37% | 134.07±68.51% | 123.95±58.39% | 140.22±76.00% |
| sham-tDCS | GABRA3 rs4828696<br>‘Variant Expression’ | 54.89±61.47% | 59.22±38.61% | 56.78±32.96% | 131.35±98.61% | 160.24±130.49% | 141.79±80.21% |
| | GABRA3 rs4828696<br>‘Normal Expression’ | 42.54±35.78% | 50.47±43.61% | 61.84±71.58% | 113.94±60.60% | 115.01±64.83% | 132.56±84.79% |

## Discussion

This study investigated response variability to a common a-tDCS protocol and whether the identified inter-individual variability was in-part genetically mediated. It was hypothesised that naturally occurring variations in genes that encode for NMDA and GABA receptors involved in excitatory and inhibitory cortical pathways would influence inter-individual variability to a-tDCS. Subgrouping subjects via pre-determined post a-tDCS response thresholds and statistical cluster analyses identified two distinct groups of individuals significantly different from one another. A group of subjects that responded with significant increases in CSE (i.e. responders) and another that responded with no increase or reductions in CSE (i.e. non-responders) were identified. No significant changes in ICI or ICF were observed in both responder and non-responder subgroups.

No significant relationships were identified between a-tDCS response and the expression of NMDA receptor genes. Significant relationships were however identified for GABA receptor gene expression, revealing that subjects with a variation in the expression of the GABRA3 gene, which encodes for inhibitory GABA-A receptors, were significantly more likely to be categorised as a-tDCS responders when compared to those with normal GABRA3 gene expression.

### Group and subgroup level analysis

To these author’s knowledge, this current study with a sample size of sixty-one healthy male volunteers is the largest of its kind that has investigated inter-individual variability in response to a-tDCS. In addition to reporting overall significant increases in CSE following a-tDCS, the reported subgroups of individuals based on subjects exceeded a pre-determined threshold was similar to previous studies. Based on the pre-determined threshold of sham-tDCS baseline MEP SD (Ammann et al., 2017), 43% of individuals responded as expected with increases in CSE, similar to that of previous subgrouping studies ranging from 33-74% (Ammann et al., 2017; Chew et al., 2015; López-Alonso et al., 2014, 2015; Puri et al., 2015, 2016; Strube et al., 2015, 2016; Tremblay et al., 2016; Wiethoff et al., 2014). As a point of difference to the majority of previous large-scale subgrouping studies, by setting the threshold at the SD of CSE at baseline in the sham-tDCS condition, this subgrouping technique in-part accounted for the inherent variation that may occur when measuring changes in CSE via TMS-evoked MEPs and that is specific to the current cohort of participating individuals. If the inherent variation, as indicated by the SD in baseline conditions, is greater than 10% or 20%, as with this current study, and it is not accounted for, then it is unclear whether categorisation as a responder based on a 10% or 20% threshold as previously utilised (Chew et al., 2015; López-Alonso et al., 2014, 2015; Puri et al., 2015, 2016; Strube et al., 2016; Wiethoff et al., 2014) reflects a true increase in CSE or just the inherent variability of TMS-evoked MEPs. Failure to account for this may therefore lead to incorrect categorisations as a-tDCS responders.

To support these subgrouping findings, this current study conducted statistical cluster analyses on the largest sample-size within the tDCS literature to-date. Similar to 43% as above, a distinct cluster of 38% (n=23) of individuals were identified as responding with increases in CSE (figure 4). In comparison to previous similar studies, the percentage of individuals clustered as increasing CSE was slightly smaller. Of the seven previous studies to conduct cluster analyses, two studies did not report distinct clusters of individuals (Chew et al., 2015; Puri et al., 2016), while five reported distinct clusters of individuals that responded with increases in CSE following a-tDCS ranging from 42-47% (López-Alonso et al., 2014, 2015; Puri et al., 2015; Strube et al., 2016; Wiethoff et al., 2014). Reasons for this marginally discrepancy in percentage of responders may lie in the differences in a-tDCS protocols. To maintain consistency with the wider tDCS literature, the current study administered a-tDCS at the common stimulation intensity and duration of 1mA and 10-minutes respectively. When compared to the current study, previous stimulation durations of 13-minutes yielded greater proportions of individuals clustered in the subgroup that responded with increases in CSE (López-Alonso et al., 2014, 2015; Strube et al., 2016). Similarly, current intensities greater than 1mA categorised more individuals in the cluster that responded with increases in CSE (Puri et al., 2015; Wiethoff et al., 2014), with previous studies reporting the number of individuals responding to a-tDCS with increases in CSE increase when the current intensity increases from 1mA to 2mA (Ammann et al., 2017; Chew et al., 2015). Our current electrode montage with a relatively smaller anodal (6×4cm) electrode size compared to the cathode (7×5cm) was an attempt to increase the current density over the anode compared to the cathode, the slightly smaller current intensity and shorter stimulus duration when compared to previous studies may have contributed to the slightly smaller proportion of individuals grouped in the cluster that reported increases in CSE. Overall however, the results of both subgrouping techniques utilised in this current study add further weight to the presence of inter-individual variability in response to a-tDCS.

The above reported changes in CSE following a-tDCS and the differences in response between a-tDCS responders and non-responders were not supported by the recorded measures of cortico-cortical excitability. This study investigated whether observed increases in CSE following a-tDCS at the group level were in-part explained by either increases in ICF or reductions in ICI, or whether these changes were amplified in the a-tDCS responder subgroup. This was to explore potential mechanisms behind discrepancies in changes in CSE between a-tDCS responders and non-responders and is the first of its kind to investigate cortico-cortical excitability and a-tDCS inter-individual variability and include a sham-tDCS condition. No significant changes were reported in ICI and ICF. These results are not consistent with previous studies with small sample-sizes reporting reductions in ICI, as measured by SICI, following tDCS (Kidgell et al., 2013; Stagg et al., 2009), but are consistent with previous large sample-size studies investigating inter-individual variability to a-tDCS, reporting no significant changes in ICI and ICF between responders and non-responders (López-Alonso et al., 2014, 2015; Strube et al., 2015). The novel inclusion of a sham-tDCS condition in this current study supported our findings further by revealing no significant differences in ICI or ICF levels between a-tDCS and sham-tDCS conditions.

With the knowledge that the level of cortical excitation and LTP-like changes can be in-part regulated by GABAergic inhibition (López-Alonso et al., 2014; Murakami et al., 2012; Michael A. Nitsche et al., 2004, 2012), the results of this current study suggest that other GABAergic inhibition mechanisms, other than GABA-A receptor activity may be responsible for explaining the differences in CSE between a-tDCS responders and non-responders. This therefore opens up future opportunities to investigate other mechanisms of cortico-cortical excitability mechanisms that may differ between responders and non-responders. These may include investigation into the role other GABA receptors involved in motor cortex plasticity such as GABA-B receptors (McDonnell et al., 2006) play in the categorisation of responders and non-responders following a-tDCS. One such future study to develop deeper insights into the explanations behind inter-individual variability to a-tDCS may be to investigate changes in long-interval intracortical inhibition, a measure of GABA-B receptor activity, on the categorisation of individuals as responders or non-responders to a-tDCS.

### Genetic analysis

To these current author’s knowledge, this is the first study within the tDCS literature to identify specific gene SNPs involved in the expression of NMDA and GABA receptors and investigate their relationship to previously observed inter-individual variability. The novel finding that two gene SNPs along the inhibitory cortical pathway (GABRA3 SNPs rs1112122 and rs4828696), rather than the excitatory cortical pathway, is associated with a-tDCS response categorisation, may suggest a relationship between GABRA3 gene variants, a reduced capacity for ICI and a-tDCS responder categorisation.

The GABRA3 gene is located on chromosome 4 and encodes for the GABA-A receptor subunit α3 (Hung et al., 2013) which is expressed in abundance throughout the cerebral cortex and important for GABAergic inhibitory regulation of the central nervous system (Sieghart & Sperk, 2002). No previous studies have investigated GABRA3 gene SNPs and inter-individual variability to a-tDCS, however looking to the pharmalogical and TMS literature there are previous reports linking the GABRA3 gene SNPs rs1112122 and rs4828696 with drug-resistance in the excitotoxic neurological disease epilepsy (Hung et al., 2013). Resistance to drugs that reduce cortical excitation suggests GABRA3 gene SNPs such as rs1112122 and rs4828696 may influence the resultant expression and function of GABA-A receptors within the central nervous system, thus influencing the capacity of GABAergic inhibitory regulation (Hung et al., 2013). The reported significant increases in CSE and significant associations between GABRA3 gene SNPs rs1112122 and rs4828696 and a-tDCS responder categorisation may therefore reflect a reduced capacity for GABAergic inhibitory regulation in the a-tDCS responders.

It is important to note however that despite the significant associations between GABRA3 gene SNPs rs1112122 and rs4828696, when accounted and controlled for all of the selected SNPs, no associations were reported for SNP rs1112122 and the association trended towards significant predictive value (p=0.073). Additionally, when investigated in isolation, while an association between GABRA3 gene SNP rs4828696 ‘variant expression’ and a-tDCS response categorisation is reported in this current study, when compared to the ‘normal expression’ group, no significant differences in CSE and cortico-cortical measures of ICI and ICF were reported.

This novel relationship may suggest a link between genetic variants in GABA receptors and a-tDCS responder categorisation. Further research into the role genetic polymorphisms play in regulating CSE changes following a-tDCS, with interactions between multiple genetic polymorphisms and their effect on CSE and cortico-cortical excitability must be conducted. To these author’s knowledge, a study of this nature within the tDCS and wider NIBS literature is yet to be conducted. Study designs that facilitate the analysis of interactions between specific genetic polymorphisms and their effects on CSE and cortico-cortical excitability as well as tDCS response rates (i.e. large sample-sizes) should therefore become a focus of future studies to further investigate the contribution of genetic variations between individuals to previously reported inter-individual variability in response to tDCS.

### Limitations

The results of this current study were only reported on healthy young males. In an attempt to minimise a potential source of inter-individual variability, females were excluded based on fluctuations in estrogen and progesterone levels influencing changes in CSE (Inghilleri et al., 2004; Smith et al., 2002; Zoghi et al., 2015). Therefore, whether these current results can be extrapolated to healthy young female populations remains unclear. It is also uncertain whether similar extrapolations can be made for healthy older populations or pathological populations or whether these findings can be generalised to all forms of tDCS or just when conventional electrode montages are utilised.

### Future Directions

With the exception of GABRA3 gene SNPs rs1112122 and rs4828696, no relationships were reported between a-tDCS response category and the selected SNPs. The results of the multivariate logistical regression analysis reporting no predictive value for any particular SNP when all other SNPs were accounted for highlights the need for further investigation into interactions between the selected genes to gain additional insight into the genetic mediation of a-tDCS response categorisation. To these author’s knowledge this current study reported findings in the largest sample-size in the tDCS literature, however a sample-size of sixty-one was not sufficient to subgroup individuals based on the presence or absence of multiple gene SNPs as the resultant subgroups of individuals were too small to conduct meaningful statistical analyses. While appreciating the difficulties in recruiting large sample-sizes, future studies should aim to build upon these current findings by continuing to recruit even larger sample-sizes that may facilitate the subgrouping of individuals based on the interactions between multiple relevant gene SNPs.

Future studies should investigate the potential relationship between GABRA3 gene SNPs rs1112122 and rs4828696 and a-tDCS responder category by investigating its interaction with other gene SNPs. For example, the presence of GABRA3 gene SNPs rs1112122 or rs4828696 combined with normal expression of genes encoding for neuroreceptors involved in excitatory cortical pathways (e.g. BDNF, GRIN1 or GRIN2B) may provide additional insight into inherent genetic expressions that contribute to an increased likelihood of a-tDCS responder categorisation.

Additionally, SNPs in other genes encoding for key regulators of cortical pathways should also be investigated. The results of this current study reporting no significant changes in SICI following a-tDCS suggest GABA receptors other than GABA-A may be involved in the observed changes in CSE following a-tDCS. Investigating SNPs in genes that encode for other GABA receptors such as GABA-B receptors may provide further insight into the genetic component of GABAergic regulation of cortical plasticity following a-tDCS and the differences between individuals who respond as expected and those who do not. Lastly, the suggestions for future studies to recruit larger sample-sizes to facilitate investigate into the interaction between multiple genes can also extend to other populations such as females, older adults and pathological population and to other tDCS protocols such as c-tDCS, to investigate the relationship between genetic variation and c-tDCS response categorisation.

More broadly, future studies investigating inter-individual variability further should recognise that not all individuals that volunteer to participate are the same. With comprehensive efforts to maximise subject homogeneity in this current study, isolation of one specific factor to investigate its role in the observed inter-individual variability was permitted. Controlling for age range, gender, time-of-day and caffeine consumption ensured the cohort of subjects was as similar as possible, thus highlighting genetic variations between individuals as the differing factors that may influence response to a-tDCS. Future studies should aim to follow similar rigorous methodological measures to systematically investigate each factor previously proposed (Li et al., 2015; Pellegrini et al., 2018a; Ridding & Ziemann, 2010) as contributing to inter-individual variability.

## Conclusions & Implications

The significance of this current study is it highlights that despite consistent tDCS stimulus parameters, not all individuals are the same and cortical plasticity, as induced by tDCS, does not occur to the same extent in each individual. The results of this current study suggest that an individual’s response to a-tDCS with increases in CSE may be more likely if gene polymorphisms are present in genes encoding for GABA-A receptors. While the results of this current study do not definitively conclude whether inter-individual variability to a-tDCS is genetically mediated, they do offer potential links with GABA-A receptor gene variations that warrant exploring in future studies.

These findings that not all individuals are the same, have implications for future research as the tDCS field progresses to more consistent and meaningful applications in neurological and pathological populations. In the clinical therapeutic setting, predictability of intervention outcome is important. With ongoing investigation into the potential factors that influence inter-individual variability to tDCS, identifying and systematically investigating specific factors or combinations of factors that may have predictive value for tDCS response are important steps forward. Specifically, knowing that there is a potential relationship between GABA-A receptor gene polymorphisms and an increased likelihood of responding to a-tDCS with increases in CSE offers an opportunity for a potential genetic screening tool for suitability for tDCS. Future studies investigating interactions between specific gene SNPs, provides potential for the identification of particular combinations of gene expression that reliably predict whether an individual will respond as expected to tDCS. This may inform the specific tDCS parameters to utilise to ensure the desired effect of increases in CSE in response to a-tDCS. The results of this current study therefore offer important insight into potential avenues moving forward in the ongoing endeavour to optimise delivery of tDCS to all individuals.

## Author Contributions

Conceived and designed study: MP, SJ, MZ; Carried out data collection: MP; Conducted statistical analysis: MP; Interpreted the findings: MP, SJ, MZ; Wrote the manuscript: MP; Writing and editing of drafts: MP, SJ, MZ.

